# Kente: A Graph-based Pangenomic Approach for Horizontal Gene Transfer Detection in Microbiomes

**DOI:** 10.64898/2026.06.22.733643

**Authors:** Natalie Kokroko, Richa Jayanti, Nicolae Sapoval, Michael G Nute, Luay Nakhleh, Todd J. Treangen

## Abstract

**Motivation:** Horizontal gene transfer (HGT) shapes bacterial evolution and microbial ecosystems, yet detecting HGT within microbiomes remains a challenge due to fragmented metagenomic assemblies, reference bias, reliance on gene boundaries, and limited ability to model structural mosaicism and patterns across genomes.

**Methods:** We present Kente, a novel pangenome graph-based framework designed for HGT detection that aligns metagenomic assembly contigs to a curated database of >600 genus-level bacterial pangenome graphs constructed using minigraph. Kente infers local taxonomic composition along contigs using alignment evidence and classifies candidate transfers using structured clade-transition topologies (e.g., A-B-A sandwich, open tips, and mosaic patterns). A complementary intra-genus module detects inter-species transfers within a single genus graph using segment-level clade annotations.

**Results:** Across simulated intra- and inter-genus transfer scenarios, Kente achieves higher precision and comparable recall relative to existing gene-centric microbiome HGT detection approaches while reducing false positives from fragmented assemblies. Application to real human gut metagenomes (HMP2, *n* = 26) demonstrates Kente’s ability to detect candidate cross-lineage transfer regions in complex microbial communities. Runtime profiling shows near-linear scaling with input size, enabling efficient analysis of large metagenomic assemblies.

**Availability and Implementation:** https://github.com/treangenlab/Kente

## 1. Introduction

Horizontal gene transfer (HGT) is a major driver of bacterial genome evolution, enabling the rapid dissemination of antibiotic resistance genes, virulence factors, and metabolic innovations across microbial communities (Treangen and Rocha, 2011). In dense ecosystems such as the human gut microbiome, HGT accelerates adaptation by facilitating genetic exchange across phylogenetically distant lineages (McInnes et al., 2020). Mechanistically, HGT occurs through transformation, conjugation, and transduction (Arnold et al., 2022).

Current bioinformatics HGT detection tools broadly fall into three categories: composition-based, phylogeny-based, and homology-based. Composition-based methods identify HGT by detecting atypical G+C content (or codon usage bias) that deviate significantly from those of the host bacteria (Lawrence and Ochman, 1997). These methods rely on the premise that transferred DNA retains the compositional characteristics of the donor species or genus, leaving a detectable signature in the recipient genome. Phylogenetic methods infer HGT by comparing gene trees to species trees, identifying incongruences where genes from distantly related taxa cluster together despite separation in the species phylogeny. In contrast, similarity- or alignment-based methods detect potential transfers by identifying genes whose closest matches occur in distant lineages.

HGT detection tools implement variations of these strategies. DarkHorse (Podell and Gaasterland, 2007) and WAAFLE (Hsu et al., 2025) fall within the similarity-based category. DarkHorse identifies phylogenetically atypical genes by performing BLAST comparisons against reference databases and computing a Lineage Probability Index (LPI) score. Low LPI scores indicate greater phylogenetic distance between a gene and its host lineage, signaling potential HGT. WAAFLE detects HGT in metagenomic assemblies by aligning Open Reading Frames to a taxonomically annotated gene database and determining whether the gene content of each contig (a contiguous assembled DNA sequence reconstructed from metagenomic reads) is best explained by one or multiple clades. MetaCHIP (Song et al., 2019) adopts a hybrid framework, first identifying candidate HGT events through group-based BLAST comparisons and subsequently validating them through gene-species tree incongruence.

Read-mapping–based approaches such as Daisy and DaisyGPS (Trappe et al., 2016; Seiler et al., 2019) detect horizontal gene transfer events by leveraging split-read and paired-end mapping signatures across reference genomes. By identifying sequencing reads that align to multiple genomic locations, these methods infer candidate transfer regions with breakpoint-level resolution, achieving base-pair precision without reliance on gene annotation. However, such approaches operate on raw sequencing reads and depend on the presence of both donor and recipient reference genomes. As a result, their applicability is limited in fragmented metagenomic assemblies, where read-level signals are lost and complete reference genomes may be unavailable.

Traditional phylogenetic approaches are sensitive to gene tree estimation errors, often producing false positives when branch support is weak (Than et al., 2008). Modern phylogenetic approaches integrate machine learning techniques and reframe the problem as an outlier detection task (Mayer et al., 2026), but still remain dependent on the quality of the starting species and gene trees. Although phylogenetic networks better model reticulate evolution, their computational complexity limits large-scale application. These limitations motivate scalable alternatives that capture structural signatures of horizontal transfer without explicit tree or network reconstruction. Despite their effectiveness, existing HGT detection approaches face certain practical and conceptual challenges. Many methods analyze genes or short sequences individually, limiting their ability to capture structural patterns across entire genomes. Similarity-based approaches often require extensive homology searches against large reference databases, which introduce significant computational bottlenecks. Furthermore, dependence on a single reference genome can lead to reference bias, as horizontally transferred regions often reside within the accessory genome and may be absent from the chosen reference (Secomandi et al., 2025).

Recent advances in graph-based genome representations offer a powerful alternative to traditional linear genome models. By integrating multiple genomes into a unified representation, these graphs inherently mitigate single-reference bias, allowing for more accurate detection and characterization of complex structural variants (Curry et al., 2024).

The concept of the pangenome captures the full complement of genetic diversity within a taxonomic lineage, comprising both a conserved core genome shared across all members and an accessory genome that varies among strains (Tettelin et al., 2005). Pangenome graphs extend this concept by representing genomic segments as nodes and connecting adjacent segments within a genome with edges (Guarracino et al., 2022). In this framework, each genome corresponds to a single path through the graph, linking its constituent segments in order. When multiple genomes share a segment but differ in their surrounding sequences, several edges may converge at that node, reflecting both shared ancestry and lineage-specific variation. Recent work has explored the use of pangenome graphs to characterize genome plasticity and regions associated with HGT. In particular, tools such as PPanGGoLiN (Gautreau et al., 2020) and its extensions panRGP (Bazin et al., 2020) and panModule (Bazin et al., 2021) use partitioned pangenome graphs to identify Regions of Genome Plasticity (RGPs). These regions often correspond to genomic islands that are frequently associated with horizontal gene transfer events. However, these approaches operate on collections of complete isolate genomes and rely on grouping genes into families, which are then classified into persistent (core), shell, and cloud categories based on their frequency across genomes. As such, they do not directly support the detection of HGT events in fragmented metagenomic assemblies, nor do they leverage sequence-to-graph alignment to infer local taxonomic transitions along contigs.

### 1.1. Contributions

However, while pangenome graphs have gained widespread adoption for several bioinformatic tasks including high-accuracy read alignment and genotyping (Eizenga et al., 2020)(Garrison et al., 2024), to the best of our knowledge, pangenome graphs have not yet been explicitly leveraged as a framework for detecting horizontal gene transfer directly from metagenomic contigs via alignment-based inference of clade transitions across genus-level graph representations. In a graph-based topology, HGT events are uniquely characterized by specific path traversals across nodes or segments associated with distinct taxonomic clades (e.g., species, genera, or families) (Colquhoun et al., 2021). To address this methodological gap, we introduce Kente, a pangenome-graph-based framework designed to detect HGT events in bacterial genomes. By mapping contigs directly to genus-level pangenome graphs and identifying structured clade transitions along graph paths, Kente leverages the topological advantages of graphs to move HGT detection beyond the limitations of linear reference models.

## 2. Methods

### 2.1. Genus-level Pangenome Graphs

Kente operates exclusively within a pangenome graph framework for detecting HGT in metagenomic datasets. To enable this, we constructed a genus-level bacterial pangenome graph database from publicly available bacterial genomes obtained from NCBI RefSeq (Goldfarb et al., 2025). To minimize assembly artifacts and fragmentation-induced noise, we restricted genome selection to assemblies categorized as “Complete Genome” or “Chromosome”. Genera represented by fewer than five complete or chromosome assembly level genomes were excluded from graph construction. This threshold was chosen to ensure sufficient intra-genus diversity for distinguishing conserved core segments from accessory variation within each graph.

To represent the structural diversity within each eligible bacterial genus, we constructed pangenome graphs using an iterative sequence-to-graph mapping strategy. For each genus, the representative backbone genome, denoted as *G*_0_, was selected to minimize reference-mapping bias. We calculated pairwise MinHash distances between all *n* candidate genomes (where *n* ≥ 5) using Mash (Ondov et al., 2016), designating the genome with the highest average nucleotide identity to the rest of the set as the initial backbone. The pangenome was then modeled as a directed multigraph *G* = (*V, E*), where *V* represents the set of nodes containing double-stranded DNA segments and *E* represents the set of directed edges indicating genomic adjacency (Garrison et al., 2024). Using `minigraph` (Li et al., 2020), the remaining *n* − 1 genomes were aligned sequentially to *G*_0_, where structural variations (SVs) exceeding a 100 bp threshold were integrated as novel nodes and edges. This iterative process yielded a graph database stored in the Graphical Fragment Assembly (GFA) format, specifically utilizing the reference-based (rGFA) convention (Li, 2019). This structured framework preserves the linear coordinate system of the *G*_0_ backbone while capturing the complex structural diversity, such as large-scale insertions and mobile genetic elements, necessary for identifying HGT-driven clade transitions.

### 2.2. Contig-to-Graph Alignment

Due to the large Kente graph database which comprises over 600 genus-level pangenome graphs, aligning every contig against the full database would be computationally prohibitive. To reduce the search space while preserving sensitivity, Kente includes a taxonomic pre-profiling step using Kraken2 (Wood et al., 2019). For each metagenomic assembled contig sample, Kraken2 is applied using RefSeq Standard database (build date: 04/2025). Resulting classifications are aggregated at the genus level to identify the taxa represented in each sample. Only those detected genera are selected as candidate graphs for subsequent contig-to-graph alignment. This pre-filtering step serves strictly as a computational optimization and does not constitute taxonomic assignment within the Kente framework. All genus labels used for HGT inference are determined exclusively through competitive graph alignment scoring. Contigs lacking confident Kraken2 classification are still retained and aligned against the full graph database to avoid excluding potential donor genera absent from the initial profile.

In practice, Kraken2 pre-filtering reduces the number of candidate genus graphs per dataset from ~600 to fewer than 30 in typical metagenomic samples, substantially reducing runtime while maintaining sensitivity to inter-genus transfer events. Following candidate genus selection, contigs are aligned independently to each selected genus-level pangenome graph using `minigraph`. To maximize sensitivity toward divergent HGT events, the `-x lr` preset is utilized. Each alignment produces a Graph Alignment Format (GAF) file describing contig-to-graph mapping including CIGAR operations, mapping quality and divergence estimate scores. At this stage, all alignment evidence is retained for downstream competitive evaluation. All retained alignments are weighted using the divergence tag (dv) reported by `minigraph`.

#### Module I: Inter-Genus HGT Detection (across graphs)

Since each genus-level graph is constructed independently and maintains its own segment name space, path-based transitions cannot be compared directly across different genera. Therefore, inter-genus HGT detection is performed exclusively in query-coordinate space. Contigs were partitioned into fixed-size windows (which can be adjusted based on input data type, default: 500 bp). For each window, cumulative alignment support was computed across all candidate genera by summing the alignment support values derived from the minigraph mappings within that window. A window was assigned to the genus with the maximum cumulative support if that genus uniquely achieved the highest score among all candidate genera. If there was a tie in terms of alignment scores and divergence, both genera are called and assigned that window. Finally, if no genus claimed that window, it is assigned unlabeled. Adjacent windows sharing identical genus labels were merged into contiguous genus blocks.

For each query contig *q*, the genus *A* with the largest total alignment support (number of aligned nucleotides) across the alignment path *P*_*q*_ was assigned as the host genus of *q*. This avoids requiring a known host *a priori* and supports metagenomic assemblies where the host organism may be unknown. Given the host *A*, any contiguous segment or block *B* in the alignment path was identified as a candidate foreign insertion if it satisfied the following criteria:

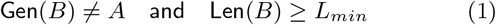

where Gen(*B*) is the consensus taxonomic label of the nodes in block *B*, and *L*_*min*_ is a minimum length threshold (default: 500 bp). This length threshold is flexible depending on the dataset the user is analyzing.

To ensure high specificity, we prioritized the detection of *sandwich topologies*, defined by the sequence *A* → *B* → *A*. In this model, a foreign block *B* is identified as a high-confidence HGT event if it is flanked upstream and downstream by host-labeled blocks (*A*) within a maximal bounded distance *λ* (default: 2 kb). This parameter *λ* allows the framework to bridge short unlabeled gaps or assembly artifacts while maintaining topological continuity.

In addition to the *A* → *B* → *A* core detections, we identified *tip insertions* (patterns of *A* → *B* or *B* → *A*). While recorded, these events were assigned a lower confidence score to account for potential artifacts resulting from contig fragmentation or incomplete pangenome representation at the sequence termini.

#### Module II: Intra-Genus / Inter-species HGT detection (within a graph)

Within a single genus-level pangenome graph, segment identifiers are shared across all embedded genome paths. This enables direct path-aware clade labeling at the segment level. For intra-genus analysis, contigs were aligned to a single genus graph. Instead of competitive cross-graph scoring, we leveraged segment-level clade annotations embedded within the graph. Each graph segment was labeled according to the species of the genomes traversing it. For graph segments shared across multiple species, all corresponding species labels were retained. Minigraph determines the highest-scoring contig traversal path through the graph. Regions with comparable support for multiple species were retained as multi-species assignments and reported as mosaic transitions rather than collapsed into a single-species label. Using the extracted graph paths from the GAF alignments, we identify transitions between segment-associated clades along the contig traversal path.

### 2.3. HGT Event Calling and A-B-A Pattern Detection

Each candidate block is classified according to its left and right flanking context along the contig. Once candidate foreign blocks are identified, Kente classifies each event based on its flanking context along the query path. Let *A* denote the inferred host taxon and *B* represent a candidate foreign block. We define *L* and *R* as the nearest taxonomically labeled flanking blocks to the left (upstream) and right (downstream) of *B*, respectively.

- **High-Confidence Sandwich Events (***A* → *B* → *A*): A candidate block *B* is classified as a high-confidence HGT event if it satisfies the condition:

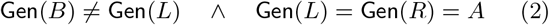

subject to the bounded distance constraint:

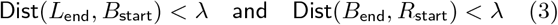

where *λ* (default: 2 kb) allows for the bridging of short unlabeled nodes, assembly gaps, or non-informative sequence segments while preserving structural continuity.
- **Tip Insertions (***A* → *B* **or** *B* → *A*): These patterns occur when a foreign segment is localized near the end of a sequence. While recorded, these are assigned lower confidence as they may represent contig fragmentation or incomplete graph representation rather than proven genomic integration.
- **Mosaic and Complex Transitions:** Paths involving multiple adjacent foreign blocks or nested taxonomic shifts are reported as complex events. These are classified separately to preserve the distinction between simple insertion events and larger, potentially multi-step recombination sites.

### 2.4. Output Data Format

For each identified horizontal gene transfer event, Kente generates a structured record in tab-separated value format to facilitate downstream statistical analysis and benchmarking. Each record represents a single foreign block and contains essential metadata including the contig identifier (`contig_ID`), total contig length (`contig_len`), the inferred host taxon (`Clade_A`), and the putative donor taxon (`Clade_B`). To define the physical boundaries of the transfer, the output specifies the genomic coordinates (`start`, `end`) and the total length of the foreign segment, alongside its structural classification (`event_type`), distinguishing between *Sandwich* (*A* → *B* → *A*), *Tip*, and *Complex* topologies. In cases where multiple structurally distinct foreign blocks are present within the same sequence, Kente reports these as independent events, thereby providing a comprehensive map of genetic mosaicism across the metagenomic assembly.

### 2.5. Experimental Evaluation Design

#### 2.5.1. Simulation-Based Benchmarking with HgtSIM

To evaluate the precision and recall of Kente, we generated synthetic horizontal gene transfer events using HgtSIM (Song et al., 2017). Simulations were designed to assess performance across (i) inter-genus transfers, (ii) intra-genus/inter-species transfers, and (iii) varying contig-length regimes that mimic fragmented and near complete genomes. For each simulation, a recipient genome was selected from the target lineage (host background), and a donor genome was selected either from a distinct genus (inter-genus experiments) or from a different species within the same genus (intra-genus/inter-species experiments).

#### 2.5.2. Inter-genus simulations

For inter-genus benchmarking, we evaluated Kente under two transfer scenarios representing different levels of taxonomic divergence. First, we selected two phylogenetically distinct genera, *Acinetobacter* and *Pseudomonas*. An *Acinetobacter* genome was designated as the recipient (host), and coding sequences from a *Pseudomonas* donor genome were inserted using HgtSIM. Second, to assess performance under a more closely related inter-genus setting, we simulated transfers between *Bacteroides* and *Parabacteroides*, two gut-associated genera within the order *Bacteroidales*. This additional benchmark was included to evaluate whether Kente remains effective when donor and recipient taxa share greater sequence similarity and more overlapping genomic content. For each inter-genus simulation, a total of 80 HGT events were simulated per recipient genome under a mixed mutation model with 5– 10% mutation rate to mimic imperfect metagenomic samples. Following genome-level simulation, modified genomes were fragmented into synthetic contigs across three varying lengths; (i) Short/fragmented contigs 1-30kb, (ii) 30-100 kb and (iii) near-complete. For each simulation we generated 80 positive contigs and 80 negative contigs. Negative contigs served as controls for evaluating false positive rates.

Because MetaCHIP requires both donor and recipient genomes to be present as independent genomes within the analyzed community, we generated an additional MetaCHIP-compatible benchmark using the *Parabacteroides*-to-*Bacteroides* simulation. Three independently seeded HgtSIM recipient assemblies were retained as MAG-like draft genomes rather than fragmented into individual benchmarking contigs, and the corresponding donor genomes were included as separate community members. MetaCHIP was then executed using its standard best-match (BM) and phylogenetic reconciliation workflow to evaluate recovery of simulated donor-recipient transfer signals under genome-centric conditions.

#### 2.5.3. Intra-genus (different species simulations)

To evaluate intra-genus detection, we simulated inter-species transfers within the *Bacteroides* genus. Specifically *B. fragilis* was selected as the recipient genome, and coding sequences from *B. ovatus* were used as donors. Because inter-species detection within a genus occurs against a highly similar background, we evaluated a broader divergence range 5–25% to stress-test robustness across increasing sequence divergence. Followed by the fragmentation into the same varying contig-lengths as the inter-genus simulations.

## 3. Results

### 3.1. Inter-genus Simulated data

Based on the simulation setup, Kente’s performance was assessed using contig and event-level precision, recall, and F1 score relative to HgtSIM’s ground truth across varying contig lengths; Table 1. Values are reported as mean ± standard error across independent simulation replicates. In the shortest contig regime (1–30 kb), Kente achieved higher precision than WAAFLE (0.774 ± 0.075 vs. 0.656 ± 0.032), but lower recall (0.188 ± 0.032 vs. 0.225 ± 0.013), resulting in a lower F1-score (0.297 ± 0.037 vs. 0.333 ± 0.012). This is consistent with WAAFLE’s stronger performance on shorter contigs and suggests that Kente is more conservative under limited sequence context. As contig length increased, Kente’s performance improved. In the 30–100 kb regime, Kente slightly outperformed WAAFLE across all three metrics, with precision, recall, and F1-score of 0.588 ± 0.007, 0.513 ± 0.019, and 0.548 ± 0.014, respectively, compared with 0.558 ± 0.027, 0.496 ± 0.036, and 0.524 ± 0.030 for WAAFLE. On near-complete genomes, Kente achieved perfect precision across replicates (1.000 ± 0.000), together with a recall of 0.579 ± 0.042 and an F1-score of 0.732 ± 0.033. This improvement reflects availability of full flanking context required for robust A-B-A topology detection for Kente. These observations match results reported in (Hsu et al., 2025), with WAAFLE performing better on fragmented contigs.

**Table 1.** Performance comparison of Kente and WAAFLE in detecting inter-genus HGT events across varying contig lengths. Values are reported as mean ± standard error across 3 independent simulation replicates. Near complete indicates contig lengths >100 kb. * indicates no detected HGT events.

| Contig Length | Tool | Precision | Recall | F1-Score |
| --- | --- | --- | --- | --- |
| 1–30 kb | Kente | 0.774 ± 0.075 | 0.188 ± 0.032 | 0.297 ± 0.037 |
| | WAAFL | 0.656 ± 0.032 | 0.225 ± 0.013 | 0.333 ± 0.012 |
| 30–100 kb | Kente | 0.588 ± 0.007 | 0.513 ± 0.019 | 0.548 ± 0.014 |
| | WAAFL | 0.558 ± 0.027 | 0.496 ± 0.036 | 0.524 ± 0.030 |
| Near Complete | Kente | 1.000±0.000 | 0.579±0.042 | 0.732±0.033 |
|  | WAAFL | * | * | * |

In the MetaCHIP-compatible MAG-like simulation, MetaCHIP identified 68 candidate HGT relationships using its best-match (BM) strategy. Following phylogenetic reconciliation with Ranger-DTL, 12 directional HGT predictions were retained. Most retained predictions correctly inferred transfer direction from *Parabacteroides* donor genomes into *Bacteroides* recipient assemblies, demonstrating successful recovery of simulated donor-recipient transfer signals when donor genomes were explicitly represented within the analyzed community. Because MetaCHIP reports genome/gene-family-level transfer relationships rather than contig-localized HGT intervals, we treated this analysis as a complementary genome-bin benchmark rather than a direct contig-level precision/recall comparison with Kente.

To evaluate performance under a more closely related inter-genus setting, we simulated transfers between *Parabacteroides* donor genomes and *Bacteroides* recipient genomes. Despite the increased sequence similarity between donor and recipient taxa, Kente maintained strong precision across all contig-length regimes and achieved perfect recall on near-complete assemblies (Table 2). Performance improved with increasing contig length, with recall increasing from 0.242±0.018 on 1–30 kb contigs to 1.000±0.000 on near-complete assemblies. In contrast, WAAFLE detected substantially fewer simulated transfer events across all contig-length regimes, resulting in markedly lower recall and F1-scores (Supplementary Table 2). These results suggest that Kente remains effective in distinguishing transfer boundaries even when donor and recipient genomes share substantial sequence similarity.

**Table 2.** Kente performance metrics for inter-genus HGT detection between closely related *Bacteroides* and *Parabacteroides* genomes across varying contig sizes. Values are reported as mean±SE across 3 independent simulation replicates. Near complete indicates contig lengths >100 kb.

| Contig Length | Precision | Recall | F1-Score |
| --- | --- | --- | --- |
| Near Complete | 0.685±0.030 | 1.000±0.000 | 0.812±0.021 |
| 30–100 kb | 0.584±0.026 | 0.633±0.023 | 0.607±0.020 |
| 1–30 kb | 0.738±0.071 | 0.242±0.018 | 0.364±0.029 |

**Table 3.** Kente performance metrics for intra-genus HGT detection across varying contig sizes. Values are reported as mean±SE across 3 independent simulation replicates. Near complete indicates contig lengths >100 kb.

| Contig Length | Precision | Recall | F1-Score |
| --- | --- | --- | --- |
| Near Complete | 1.000±0.000 | 0.950±0.026 | 0.974±0.014 |
| 30–100 kb | 0.604±0.030 | 0.723±0.023 | 0.658±0.028 |
| 1–30 kb | 0.714±0.029 | 0.473±0.024 | 0.567±0.012 |

### 3.2. Intra-genus simulated data

We next evaluated Kente’s performance in identifying transfers between closely related species within the same genus, which is a major challenge in metagenomic HGT detection. Under the intra-genus simulation within *Bacteroides*, Kente achieved high detection performance. For near-complete contigs, Kente reached an average recall of 0.950 and a precision of 1.00, performance decreased for shorter fragmented contigs with precision of 0.604 and recall of 0.723, consistent with reduced structural context in fragmented assemblies. While Kente’s graph-based algorithm accurately detected structural boundaries corresponding to inter-species transfers, species-level assignments occasionally lacked full resolution. In some cases, donor or recipient segments were assigned to a broader genus-level label (e.g. *Bacteroides*) or to a closely related sister species. Importantly, Kente consistently identified the presence of a transfer event between genomic lineages within the genus. Under the same simulation conditions, other evaluated tools (WAAFLE and MetaCHIP) did not report any candidate intra-genus HGT events.

### 3.3. Real Data Evaluation on HMP2

To evaluate Kente on real metagenomic data, we analyzed 26 assembled stool metagenomes from control participants in the HMP2 (IBDMDB) (Lloyd-Price et al., 2019); we leveraged the publicly available assemblies used in (Hsu et al., 2025). We applied both WAAFLE and Kente to the same assembled contigs without additional preprocessing or reassembly. Overlap between the predicted HGT events is summarized in Figure 2a. Under strict structural criteria (A-B-A only), Kente identifies 1,607 candidate HGT events, whereas WAAFLE predicts 598 events, with only a small subset shared between the tools. When Kente detection was expanded to include tip (A-B/B-A) and mosaic configurations, total calls increased to over 10,000 putative HGT events, yet the overlap with WAAFLE remained limited (increased from <1% to ~20% same candidate HGT contigs). This modest intersection suggests that the two methods capture distinct classes of candidate transfers. Based on this, we further analyzed both tool results and found contig-level distributions could account for the major difference in the candidate HGT calls. As displayed in Figure 2c, WAAFLE calls were concentrated in shorter contigs, with an average contig length of 4 kb. In contrast, strict A-B-A calls from Kente were enriched in substantially longer contigs with a mean length of 40 kb. These large assembled metagenomic contigs were abundant, with over 20,000 contigs per sample. When Kente detection was expanded to include AB and more complex mosaic configurations, the average contig length decreased to ~20 kb, reflecting increased sensitivity to fragmented assemblies. Together, these results suggest that WAAFLE and Kente operate in complementary detection regimes, with WAAFLE recovering gene-level signals from shorter contigs and Kente recovering graph-supported, structurally contextualized events from longer contigs. These patterns are consistent with methodological design. WAAFLE relies on gene-level alignment and, as mentioned in their study, is optimized for shorter assembled contigs, while Kente generally requires sufficient flanking host context on both sides of a foreign block for A-B-A calls, to reduce false positives, favoring longer contigs. Inspection of WAAFLE output revealed that a substantial fraction of its calls correspond to open A-B patterns rather than the fully flanked A-B-A events. In contrast, Kente distinguishes between high-confidence “sandwich” HGT events and open/mosaic configurations. Despite methodological differences, both tools predominantly identified transfers occurring within closely related genera common to the human gut microbiome, particularly *Bacteroides* and *Parabacteroides*.

**Figure 1.**
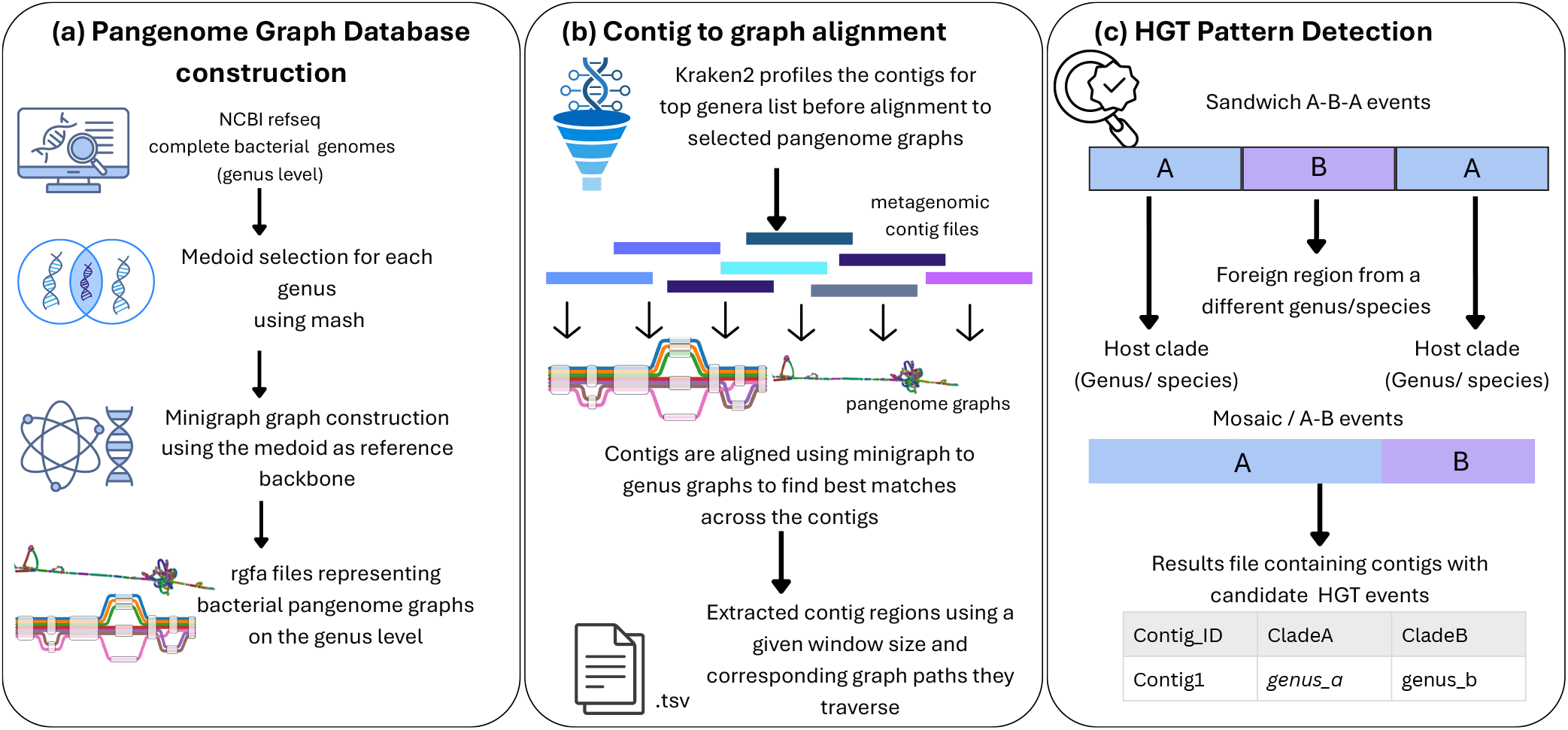
Overview of Kente. (a.) A pangenome graph database was pre-built using `minigraph` which contains genus level bacterial pangenome graphs. (b.) Kente aligns contigs to the graph database to find best matches across different regions of contigs. (c.) These alignments are extracted per contig region to find distinct A-B-A/A-B patterns along contigs which signify potential HGT events.

**Figure 2.**
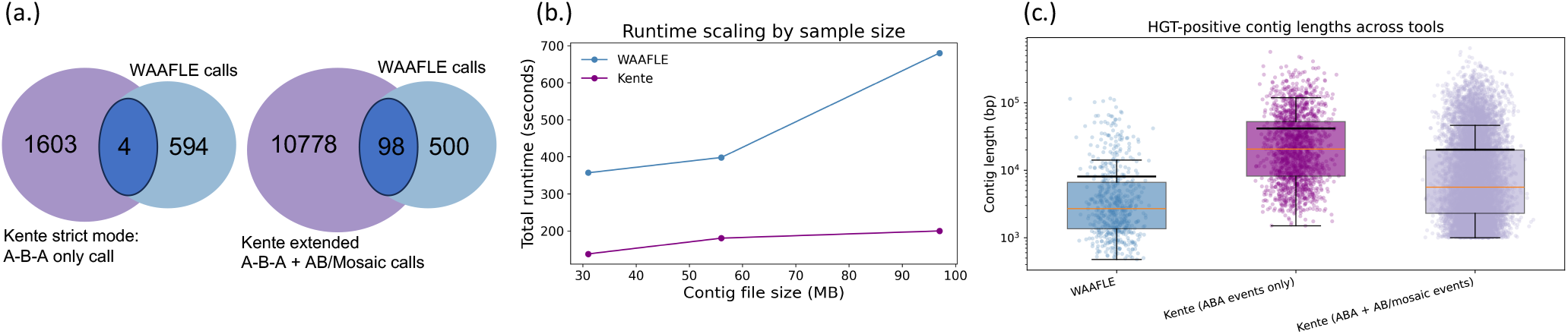
(a.) Comparison of HGT calls between Kente and WAAFLE on HMP2 stool metagenomes (b.) Runtime (seconds) across datasets of increasing size for each tool. (c.) Distribution of contig lengths for HGT-positive contigs across tools

#### 3.3.1. Taxonomic Distribution of HGT in Stool

Across the 26 HMP2 stool metagenomes, detected HGT events were predominantly observed among gut-associated genera within the Bacteroidetes lineage. For Kente, the most frequent transfer pairs as observed in Figure 3b involved *Parabacteroides, Phocaeicola*, and *Bacteroides*, with recurrent exchanges observed between *Parabacteroides* ↔ *Phocaeicola* and *Phocaeicola* ↔ *Bacteroides*. However, WAAFLE recovered a more limited subset of these relationships. Notably, the high volume of transfers between *Phocaeicola* and *Bacteroides* is consistent with their shared taxonomic history as many highly abundant gut species now assigned to *Phocaeicola* were recently reclassified from the genus *Bacteroides* (García-López et al., 2019). Despite these differences, both methods identified comparable patterns involving recurrent *Alistipes*-associated transfers and frequent associations with *Bacteroides*-related lineages. In particular, both approaches identified 23 HGT interactions between *Alistipes* and *Bacteroides*, providing evidence of concordance between the two methods despite their differing detection strategies. Together, these observations suggest that taxonomic representation and lineage resolution may influence the transfer profiles recovered by different approaches.

**Figure 3.**
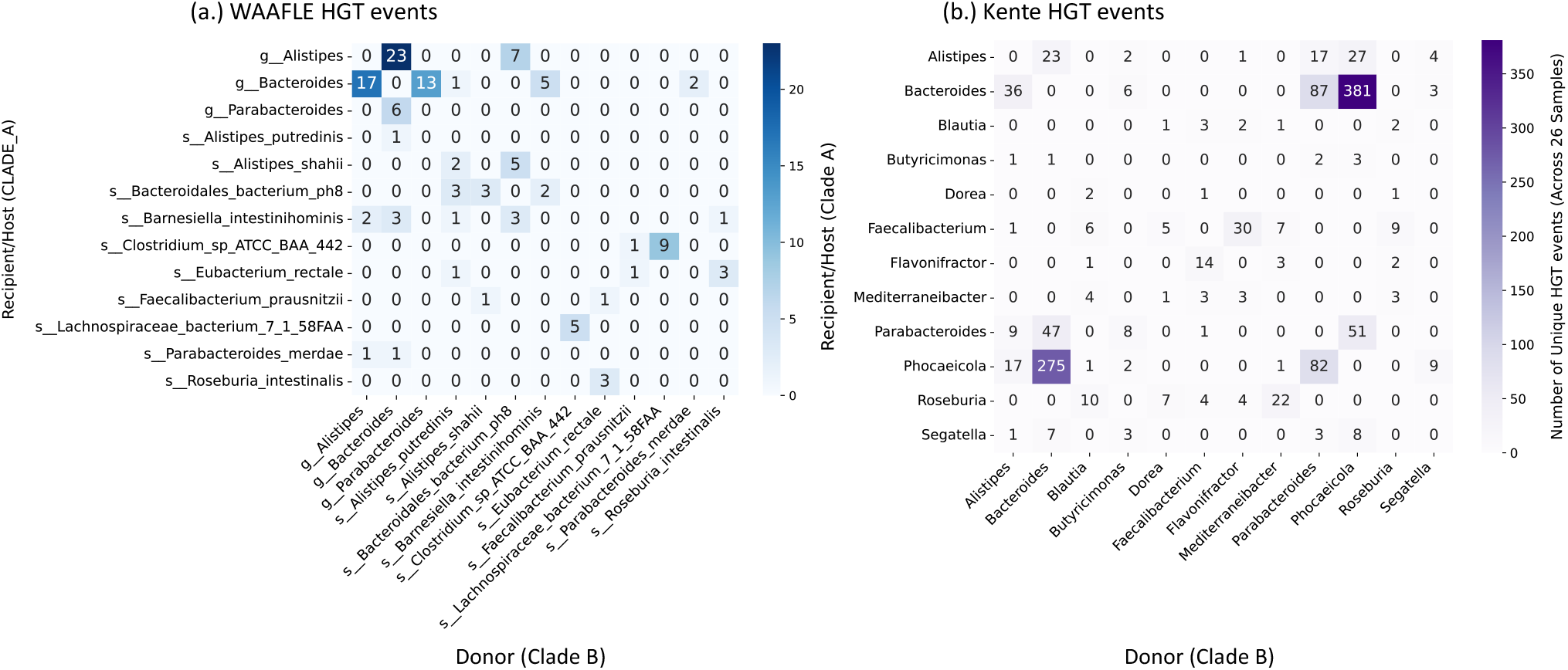
Taxonomic distribution of predicted HGT events in 26 HMP2 stool metagenomes. Heatmaps show donor–recipient transfer pairs detected by (a.) WAAFLE and (b.) Kente across the same HMP2 assembled stool samples. Rows indicate the inferred recipient or host clade, and columns indicate the inferred donor clade. Cell values represent the number of unique predicted HGT events across the 26 samples.

### 3.4. Computational Efficiency

Table 4 reports the peak memory usage of Kente. Figure 2b indicates runtime requirements of Kente on different input file sizes (from 30MB assembled metagenomic sample to over a 100MB fasta files). Kente takes less time, even with larger metagenomic files. All software analysis was performed on a Linux system, with 16 threads allocated per run for both WAAFLE and Kente. The (`usr/bin/time`) command was used to report the time and memory statistics. To assess the cost of removing the taxonomic pre-filtering step, we performed an additional test mapping a single HMP2 stool assembly against the full genus pangenome graph database. Sequential mapping across all graphs required approximately 14.8 minutes of wall-clock time and ~ 3.25 GB of peak RSS. While this mode avoids the memory overhead of kraken2, it increases the total Kente runtime from about 3 minutes to 20 minutes per sample.

**Table 4.** Peak memory usage and runtime comparison between Kente and WAAFLE across the HMP2 stool assembled dataset.

| Tool | Memory (GB) | Runtime (min) |
| --- | --- | --- |
| Kente (without Kraken2) | 3.2 | 20.0 |
| Kente + Kraken2 | 87.0 | 2.2 |
| WAAFLÉ | 8.6 | 6.0 |

Memory usage for Kente + Kraken2 includes loading the full Kraken2 standard database into RAM (approximately 84 GB), while the Kente graph-alignment framework itself contributes approximately 3 GB peak RSS.

## 4. Discussion

In this study, we introduce Kente, a pangenome-graph-based framework for detecting horizontal gene transfer (HGT) in metagenomic assemblies. By leveraging genus-level pangenome graphs in rGFA format constructed with `minigraph`, Kente reframes HGT detection from a similarity-based, gene-centric problem to a taxonomy discordance-based, genome-graph-centric problem. Most pangenome analyses and HGT detection methods remain gene-centric. However, horizontal gene transfer can occur at sub-gene resolution through partial gene exchange, recombination, and mosaicism (Chan et al., 2011). Prior efforts, such as tMHG-Finder, have been proposed to address sub-gene mosaicism within gene-centric frameworks (Yin et al., 2025). These approaches represent an important step toward recognizing gene-fragment level recombination events within bacterial communities. However, modeling sub-gene mosaicism within all-vs-all gene-centric blast-based frameworks, even when accelerated with guide trees, becomes increasingly computationally complex as the number of genomes scales to tens of thousands of genomes.

By constructing genus-level pangenome graphs in rGFA format, Kente represents both core and accessory genomic diversity within a unified coordinate system. Each genome corresponds to a path through the graph, and structural variants are encoded as alternative nodes and edges. This representation further reduces the dependence on a single reference. Our results demonstrate that Kente’s graph-based approach enables improved HGT detection across a range of phylogenetic distances. In inter-genus simulations, performance improved with increasing contig length, reflecting the importance of flanking structural context for high confidence A-B-A detection. In intra-genus simulations, Kente maintained a strong precision and recall even when donor and recipient species are highly similar. Applying Kente to the 26 HMP2 stool metagenomes revealed transfer patterns concentrated among prevalent gut-associated genera (Figure 3). Notably, the limited overlap observed between Kente and WAAFLE predictions should not be interpreted as disagreement between methods, but rather as evidence that graph-based and gene-centric frameworks capture partially distinct components of the HGT signal. WAAFLE is well suited to detecting gene-level discordance in shorter assembled contigs as shown in Figure 2c, while Kente benefits from longer contigs that preserve sufficient flanking context for resolving structured clade transitions along graph paths. Rather than competing approaches, Kente and WAAFLE may be considered complementary, with combined application offering more comprehensive HGT discovery across metagenomic datasets characterized by heterogeneous contig-length distributions.

Despite its strengths, it is important to note that Kente’s detection sensitivity depends on representation of donor lineages within the graph database. Transfers from taxa absent in the database may remain unlabeled or incorrectly assigned to closely related genera. Expanding graph coverage or incorporating higher-level taxonomic graphs (e.g., family-level representations) may mitigate this issue. Moreover, highly fragmented assemblies limit detection of high confidence HGT events due to insufficient flanking context. Finally, although rGFA preserves coordinate structure relative to a backbone genome as designed by `minigraph`, complex structural rearrangements may be simplified within graph construction. Future extensions of Kente will explore alternative graph construction strategies to better capture complex structural variations and potentially reduce the backbone bias. In particular, approaches such as the minigraph-cactus pipeline (Hickey et al., 2023) that leverage multiple genome alignment may provide more balanced representations of genomic diversity and improve the detection of complex HGT events. In summary, to the best of our knowledge, Kente represents a first of its kind approach for computationally-efficient microbiome HGT detection via pangenome graphs, performing favorably compared to state-of-the-art existing approaches.

## 5. Acknowledgments

NK, RJ, MGN, and TJT are supported in part by NIH NIAID P01-AI152999, and NSF awards IIS-2239114, EF-2126387. NS and LN are supported in part by the NSF grants DMS/NIGMS-2153704, DBI-2030604.

## Abbreviations

HGT: 
GAF: 
rGFA: 
SV: 
MAG: 

## Notes

### Competing Interest Statement

The authors have declared no competing interest.

