## Supplementary Table 2 for "Kente: A Graph-based Pangenomic Approach for Horizontal Gene Transfer Detection in Microbiomes"

June 3, 2026

#### Overview

This supplementary document provides additional methodological details and extended results supporting the Kente framework for detecting horizontal gene transfer (HGT) in metagenomic assemblies.

#### S1. Pangenome Graph Database Construction

The graph construction software required:

- Minigraph version $0.21 - r606$
- Mash version 2.3

The following command was used to build the pangenome graphs at the genus level. Here, backbone represents the medoid chosen for each genus by Mash.

```
minigraph -cxggs -l 1000 -L 100 -t "$THREADS" "$BACKBONE" "${FILES[@]}" \
  > "$OUTDIR/${GENUS}.rgfa" 2> "$OUTDIR/build.log"
```

##### S1.1 Special case: Escherichia

The *Escherichia* genus group initially comprised approximately 5,000 complete genome assemblies, which exceeded practical limits for iterative graph construction. To address this, Mash-based clustering was performed to reduce the set to a representative 600-genome subset. Clustering was carried out by computing pairwise MinHash distances and selecting cluster medoids to ensure the retained genomes captured the phylogenetic breadth of the full set. This approach preserves representative genomic diversity while maintaining computational tractability.

#### S2. Simulation Design

Simulated datasets were generated at the inter- and intra-genus levels. Simulations were designed to assess detection performance across varying contig lengths and taxonomic distances. Software required for simulation and their version:

- HgtSIM version 1.1.0

- MetaCHIP version 1.10.13
- WAAFLE version 1.0

##### S2.1Genome Selection

For inter-genus experiments, a single recipient *Acinetobacter* genome and a single donor *Pseudomonas* genome were selected, representing phylogenetically distant genera. For the closely related inter-genus scenario, transfers were simulated between *Parabacteroides* (donor) and *Bacteroides* (recipient), two genera within the order *Bacteroidales* that share substantial sequence similarity. For intra-genus experiments, *Bacteroides fragilis* was used as the recipient and *Bacteroides ovatus* as the donor, enabling evaluation of inter-species detection within a single genus.

Using HgtSIM, horizontal gene transfers were simulated under the following configuration:

- Number of insertion events: 80 per recipient genome
- Insertion type: Coding sequences (CDS) selected from the donor genome
- Mutation model: Mixed mutation mode with 5–10% sequence divergence for inter-genus simulations; 5–25% for intra-genus simulations to stress-test performance across increasing sequence similarity

HgtSIM produces a detailed insertion report specifying the recipient contig, breakpoint position, inserted sequence identity, and mutation level. This report served as ground truth for all precision, recall, and F1 evaluations.

##### S2.2 Contig Fragmentation

Modified recipient genomes produced by HgtSIM were fragmented into synthetic contigs of varying lengths using a custom script. Fragmentation was guided by the breakpoint coordinates provided in the HgtSIM insertion report to ensure that simulated HGT regions were preserved intact within individual fragments. Fragment start positions were selected such that each inserted donor sequence remained fully contained within a single generated contig, preventing artificial splitting of HGT loci across fragment boundaries. Negative control fragments were sampled from genomic regions that did not overlap any insertion breakpoints. This design yielded balanced evaluation datasets consisting of 80 HGT-containing and 80 HGT-free contigs per simulation replicate across three contig length regimes: 1–30 kb, 30–100 kb, and near-complete (>100 kb).

#### S3. Results

##### S3.1 Validation of Kente on Complete genomes

###### S3.1.1 Functional characterization of Staphylococcus HGT events

To validate Kente's ability to detect candidate HGT regions in a complete genome, we applied Kente to complete, high-quality reference genomes. First, the *Staphylococcus aureus* USA300 FPR3757 genome. Most lineage switches identified by Kente occurred within *Staphylococcus aureus*, consistent with transfer among closely related genomes (strain-level). In addition, one event was detected between *S. aureus* and *S. epidermidis*, representing an inter-species transfer within the *Staphylococcus* genus.

Functional annotation of the predicted regions using Prokka [Seemann, 2014] revealed the presence of known antibiotic resistance determinants and mobile genetic elements. In particular, genes encoding the mupirocin resistance protein *mupA* and the macrolide resistance methyltransferase *ermC* were identified within predicted HGT blocks. Both genes are known antimicrobial resistance determinants in *Staphylococcus* [Patel et al., 2009] and are frequently associated with plasmids and other mobile elements that facilitate horizontal transfer [McNeil et al., 2011].

Table S1: Summary of the prokka report containing the Functional annotation of genes identified in the Kente-extracted HGT block from the *Staphylococcus aureus* USA300-FPR3757 genome.

| **Gene** | **Annotation (Prokka)** |
|---|---|
| *mupA* | mupirocin resistance protein |
| *ermC'* | rRNA methyltransferase |
| IS257R2 | IS6 family transposase |
| IS431R | IS6 family transposase |
| *topB* | DNA topoisomerase III |
| *ssb* | single-stranded DNA-binding protein |

### S4. Extended Benchmarking Results

#### S4.1 WAAFLE Performance on *Bacteroides*/ *Parabacteroides* Inter-genus Simulations

Table S2 extends the benchmarking results presented in Table 2 of the main text by including WAAFLE's performance on the closely related *Bacteroides*/*Parabacteroides* inter-genus simulation scenario. This comparison was omitted from the main text for brevity but is included here for completeness. Across all contig-length regimes, WAAFLE achieved markedly lower recall and F1-scores compared to Kente, with near-zero recall on short and medium-length contigs. On near-complete assemblies, WAAFLE detected no transfer events. These results suggest that WAAFLE's gene-level alignment strategy is less effective at resolving transfer boundaries when donor and recipient taxa share high sequence similarity, whereas Kente's graph-based topology detection remains sensitive under these conditions.

Table S2: Performance comparison of Kente and WAAFLE for inter-genus HGT detection between closely related *Bacteroides* and *Parabacteroides* genomes across varying contig sizes. Values are reported as mean±SE across 3 independent simulation replicates. Near complete indicates contig lengths >100 kb. * indicates no detected HGT events.

| **Contig Length** | **Tool** | **Precision** | **Recall** | **F1-Score** |
|---|---|---|---|---|
| 1–30 kb | Kente | 0.738±0.071 | 0.242±0.018 | 0.364±0.029 |
| | WAAFLE | 0.389±0.200 | 0.013±0.007 | 0.024±0.01 |
| 30–100 kb | Kente | 0.584±0.026 | 0.633±0.023 | 0.607±0.020 |
| | WAAFLE | 0.661±0.053 | 0.063±0.000 | 0.114±0.001 |
| Near Complete | Kente | 0.685±0.030 | 1.000±0.000 | 0.812±0.021 |
| | WAAFLE | * | * | * |
